# New mechanism-based inhibitors of aspartate transcarbamoylase for anticancer drug development

**DOI:** 10.1101/662718

**Authors:** Zhen Lei, Nan Wang, Biying Wang, Zhifang Lu, Hongwei Tan, Jimin Zheng, Zongchao Jia

**Author notes:** To whom correspondence should be addressed.; (Jimin Zheng), (Zongchao Jia).

## Abstract

Aspartate transcarbamoylase (ATCase) is a key enzyme which regulates and catalyzes the second step of *de novo* pyrimidine synthesis in all organisms. *E. coli* ATCase is a prototypic enzyme regulated by both product feedback and substrate cooperativity, whereas human ATCase is a potential anticancer target. Through structural and biochemical analyses, we revealed that R167/130’s loop region in ATCase serves as a gatekeeper for the active site, playing a new and unappreciated role in feedback regulation. Based on virtual compound screening simultaneously targeting the new regulatory region and active site of human ATCase, two compounds were identified to exhibit strong inhibition of ATCase activity, proliferation of multiple cancer cell lines, and growth of xenograft tumors. Our work has not only revealed a previously unknown regulatory region of ATCase that helps explain feedback regulation, but also successfully guided the identification of new ATCase inhibitors for anticancer drug development using a dual-targeting strategy.

## Introduction

The *de novo* pyrimidine synthesis pathway is conserved in all organisms (Evans & Guy, 2004, Jones, 1980, Lee, Kelly et al., 1985), in which the first three steps are catalyzed by carbamoyl phosphate synthetase (CPSase), aspartate transcarbamoylase (ATCase), and dihydroorotase (DHOase), respectively. CPSase initiates the pathway by catalyzing the formation of carbamoyl phosphate (CP), ATCase transits the carbamoyl of CP onto Asp to produce carbamoyl aspartate (CA), and DHOase condensates CA to dihydroorotate. Among the three enzymes, ATCase has been extensively studied, especially ecATCase-holo, which is referred as a textbook example for cooperativity effect and feedback regulation (Kantrowitz, 2012, Lipscomb & Kantrowitz, 2012) (all abbreviations related to ATCase used in this paper: ecATCase-holo for *E. coli* ATCase holoenzyme, apo-ecATCase-holo for apo form *E. coli* ATCase holoenzyme, and PALA-ecATCase-holo for PALA binding form *E. coli* ATCase holoenzyme; ecATCase for *E. coli* ATCase, apo-ecATCase for apo form *E. coli* ATCase, and PALA-ecATCase for PALA binding form *E. coli* ATCase; huATCase for human ATCase, apo-huATCase for apo form human ATCase, and PALA-huATCase for PALA binding form human ATCase). In brief, ecATCase-holo is comprised of 2 catalytic trimers and 3 regulatory dimers, and it can adopt two different states at quaternary level: a low activity and low-affinity tense state (T state) and high activity and high affinity relax state (R state). High concentration of the second substrate, Asp, triggers a domain closure of ATCase which subsequently facilitates the transition from T to R state, termed cooperativity effect (Howlett & Schachman, 1977, Krause, Volz et al., 1987). The regulatory subunits can bind different nucleotides, causing a positive or negative effect on the activity of ecATCase-holo, termed feedback regulation (Gerhart & Pardee, 1962, Wild, Loughrey-Chen et al., 1989). Differently from ecATCase which is encoded separately and functions indenpently, huATCase is fused into CAD with CPSase and DHOase, but it exhibits high conservation among primary, secondary, and tertiary structures with ecATCase (Ruiz-Ramos, Velazquez-Campoy et al., 2016). Additionally, feedback regulation and cooperativity effect are also believed to exist in CAD (Moreno-Morcillo, Grande-Garcia et al., 2017, Serre, Penverne et al., 2004).

The feedback regulation of ATCase is an important means that helps organisms balance the levels of pyrimidines and purines in cells. CTP and UTP, the end products of *de novo* pyrimidine synthesis pathway, inhibit the activity of ATCase, whereas ATP and GTP promote it. For ecATCase-holo, the binding of pyrimidines or purines not only influences the V_max_, but also causes a pronounced change of K_m_ (Cockrell, Zheng et al., 2013). In other words, pyrimidines or purines change the difficulty level for ecATCase-holo to transit from T to R state. Nevertheless, it is yet to be elucidated how pyrimidines and purines exert their effects because they bind at a position far away from the active site and ATCase structures bound with pyrimidines or purines do not show obvious differences. For ecATCase-holo, the distance between the binding position and the active site is ∼60 Å. In the case of CAD, although the exact distance remains unknown due to the lack of CAD structure, the distance would also be very long because effectors are considered to bind with CPSase of CAD (Serre et al., 2004), which is far away from the active site of ATCase (Moreno-Morcillo et al., 2017). There must be some sort of yet unknown transmission mechanism which enables the regulation.

Zooming in the active site of ATCase, many completely conserved and positively-charged residues stabilize the negatively-charged substrates, CP and Asp, including K84 from an adjacent monomer, H134, and several arginines - R54, R105, R167, and R229. Among these arginines, R167 is located at the substrate entrance point or gate of the active site. In most ATCase structures, R167 faces inward toward the active pocket (which we call R167 “in” state), whereas a handful of ATCase structures show that R167 side chain protrudes away and is positioned outside the active site pocket (which we call R167 “out” state). R167 “in” state plays several key roles for ATCase, one of which is stabilizing the substrate and/or the intermediate product (Gouaux & Lipscomb, 1990, Gouaux, Stevens et al., 1990, Ke, Lipscomb et al., 1988). The domain closure of ATCase is also closely related with R167 “in” state, the occurrence of which relies on the formation of interactions among E50, R167, and R234 at R167 “in” state (Kantrowitz & Lipscomb, 1988, Ladjimi & Kantrowitz, 1988), and domain closure cannot occur when R167 adopts “out” state. Despite of the comprehensive realization about R167 “in” state, the R167 “out” state has seemed to be so far largely neglected and the only study reported has to do with the so-called “extreme T” state (Huang & Lipscomb, 2004). The role of R167 “out” in ATCase is another puzzle that has to be settled. Besides R167, there is a short flexible loop (residues A127 to H134, which we call 130’s loop) interacting with and stabilizing R167 “in” or “out” state, which further interacts with regulatory subunit in the case of ecATCase-holo. Apart from the known location of 130’s loop at the interface between the active site and the regulatory subunit, its role also remains completely unclear.

Due to the key role of CAD in pyrimidine synthesis, its activity is upregulated in cancer cells to accommodate the high demand for nucleotides (Aoki & Weber, 1981). Thus, huATCase of CAD is a potential target for anticancer therapy. In fact, attempts have been made to use N-phosphonacetyl-L-aspartate (PALA), an analog of the reaction intermediate of ATCase, as an anticancer drug. Unfortunately, it failed in clinical trials (Grem, King et al., 1988), although it exhibited inhibition of huATCase and the proliferation of colonic cancer cell line, and extension of mean survival time of mice (Swyryd, Seaver et al., 1974, Tsuboi, Edmunds et al., 1977). The recently solved huATCase structure provided a partial rationalization for the failure (Ruiz-Ramos et al., 2016). Briefly, in the huATCase, the domain closure of one catalytic chain caused by the binding of the first PALA affects the conformation of the other two active sites in the trimer, resulting in increasingly more difficult binding of the second and third PALA. This situation would be even more pronounced in the case of CAD. Owing to the negative cooperativity of binding, PALA can only partially inhibit the activity of huATCase. Additionally, low dose of PALA is also very likely to become an activator for huATCase when assembled in CAD, as is the case in ecATCase-holo, which would make it very difficult to control a proper PALA dosing during clinical trials. The clear disadvantage of PALA warrants seeking novel inhibition strategies and new inhibitors, which would target the apo form human ATCase (apo-huATCase) and would ideally not trigger the domain closure.

Herein we report several crystal structures of ecATCase and ecATCase-holo including a wild-type apo-ecATCase-holo, in which R167 “out” state clearly observed. This represents the first case of R167 “out” conformation in an ecATCase-holo structure in absence of any mutations or ligand binding in active site. By structural comparison and analysis, we firstly observed a region of R167/130’s loop located at the interface of active site and regulatory subunit that may play a key role in feedback regulation of ATCase. We investigated the region using various approaches including crystallography, enzymology, dynamic simulation and isothermal titration calorimetry etc., and demonstrated that R167 needs to switch between “in” and “out” state during the catalytic process of ATCase to guide the entrance of Asp and help the release of carbamoyl aspartate. In addition, the conformational change of R167 is under the regulation of 130’s loop and the latter was further affected by the regulatory subunit in the case of ecATCase-holo. Therefore, we considered that this region act as a modulator in response to the signal transmitted from nucleotides binding. This standpoint is also supported by previous literature (Eisenstein, Markby et al., 1989). Since huATCase is a potential target for anticancer drugs, we, taking advantage of the newly discovered feedback regulatory mechanism, performed a virtual compound screening simultaneously targeting both the newly found regulatory region and the active site of apo-huATCase. Two compounds from the top hit list exhibited strong inhibition of both huATCase activity and the proliferation of multiple cancer cell lines. Mice xenograft tumor experiments also yielded promising results. Our work revealed a new feedback regulatory mechanism of ATCase, which successfully guided us to obtain inhibitors of ATCase for new anticancer drugs development using a dual-targeting strategy.

## Results

### The R167 “out” structure of ecATCase-holo helps uncover a previously neglected regulatory region of ATCase

The structure of ecATCase-holo obtained here is virtually identical to other ecATCase-holo structures in T state, except for the conformation of R167 (Fig 1A and B). In the structure, R167 extends outwards of the ATCase active site, which we term R167 “out” to distinguish from R167 “in” state. By analyzing all reported ecATCase-holo structures (Appendix Table S1), we found only four other structures that adopt this R167 “out” state, two of which (PDB ID: 9ATC and 4E2F) have mutations destabilizing R state of ecATCase-holo (Guo, West et al., 2012, Ha & Allewell, 1998, Newell & Schachman, 1990) and the other two (PDB ID: 1R0C and 2AIR) bind with substrate analogs or products in an unusual way (Huang & Lipscomb, 2004, Huang & Lipscomb, 2006). Thus, the structure we report here is the first wild-type apo form *E. coli* ATCase holoenzyme (apo-ecATCase-holo, and apo-ecATCase for apo form *E. coli* ATCase) with R167 “out” state, which clearly demonstrates that ecATCase-holo can adopt R167 “out” state without the influence of other factors. Because of the close proximity and multiple interactions between 130’s loop and R167, we investigated R167 together with 130’s loop. The fact that R167 can adopt both “in” and “out” state indicates a certain degree of flexibility of this region. Considering that this region is located at the gate of active site of ATCase, we speculated that this region may play a regulatory role in the catalytic process of ATCase.

**Figure 1.**
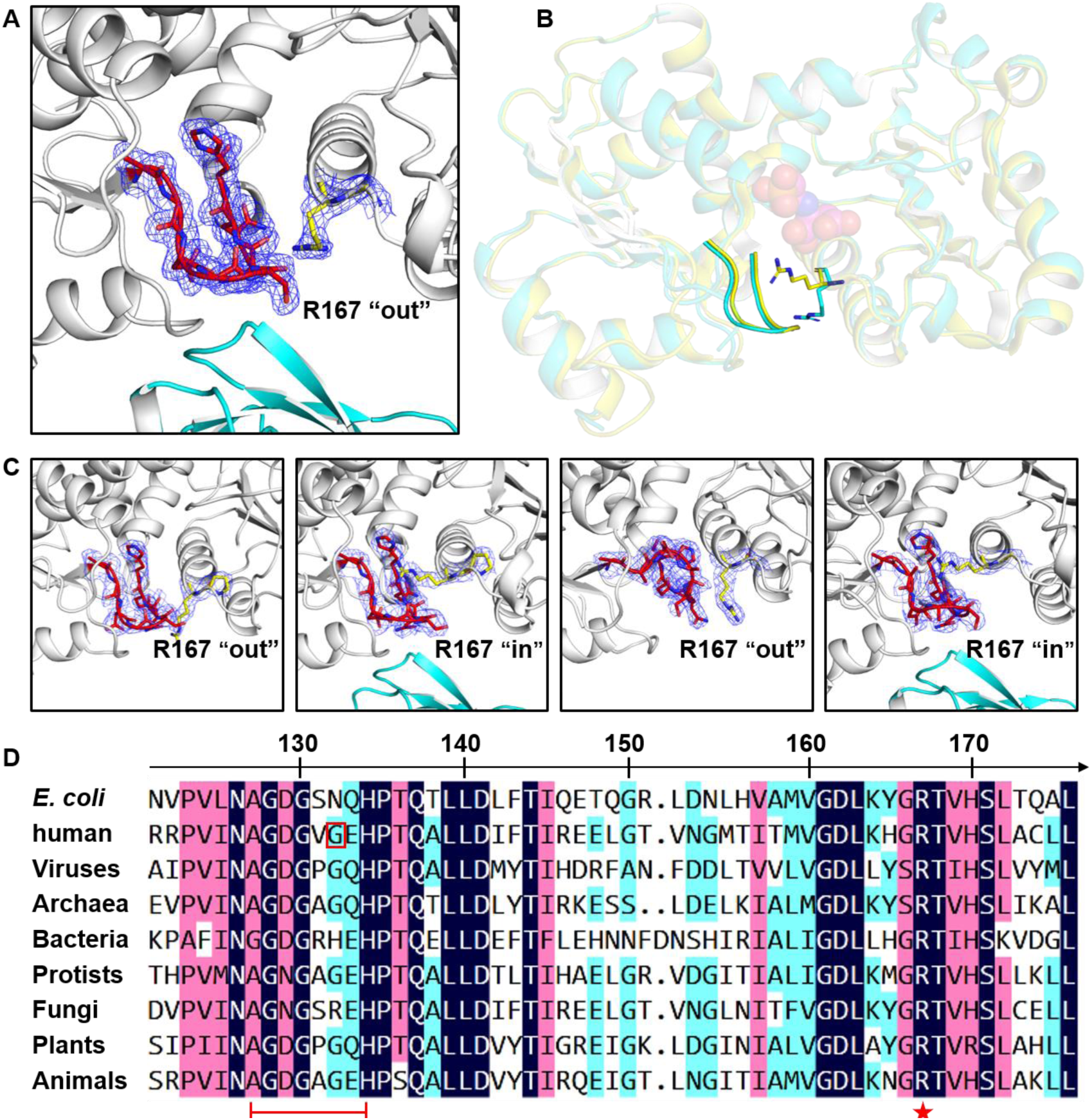
ATCase structures solved in this paper and sequences alignment of different ATCases. **A** The structure of R167/130’s loop region of wild-type apo-ecATCase-holo solved in this work, in which R167-out state is shown explicitly by electron density map (contoured at 1.0 σ). In this figure, R167/130’s loop are shown as sticks, catalytic subunit in white, regulatory subunit in cyan, R167 in red and 130’s loop in yellow. This coloring scheme is also used in other figures. **B** Comparison between the wild-type apo-ecATCase-holo structure solved in this work (cyan) and a previously reported ecATCase-holo structure (PDB ID: 1ZA1, yellow), in which R167 adopts “out” and “in” state, respectively. 130’s loop is also highlighted and the position of the active site is indicated by a docked PALA (sphere model) taken from another ATCase structure (PDB ID: 4KGV). For clarity, transparent cartoon model is used except for R167 and 130’s loop and this transparent scheme is also used in other figures. **C** Electron density maps of R167 and 130’s loop in ATCase mutants. In each graph, G166 or P166, R167 and 130’s loop are shown as sticks, and density maps were contoured at 1.0 σ. From left to right, they are G166P ecATCase, G166P ecATCase-holo, G128A/G130A ecATCase and G128A/G130A ecATCase-holo. **D** Sequence alignment of the ATCase segment containing R167 and 130’s loop in different species, from viruses to animals. R167 and 130’s loop are indicated by red star and red line, respectively. The additional glycine (G132) of huATCase is indicated by a red rectangle. See Appendix Fig S1 for the full-length alignment of selected organisms.

### Mutations that reduce the flexibility of R167/130’s loop significantly decrease the enzymatic activity of ATCase

To investigate the importance of the flexibility of R167/130’s loop, we attempted to alter local flexibility by introducing mutations and monitor their effects on enzymatic activity. G166, which is next to R167, was mutated to alanine or proline and glycines in the 130’s loop were changed to alanines, either individually or together. In this assay, ecATCase, ecATCase-holo, and huATCase were examined, and corresponding wild-type and R167A ATCase (similar mutation was previously shown to cause a dramatically decrease of ATCase activity (Stebbins, Zhang et al., 1990)) were used as positive and negative control, respectively. Our results from the aforementioned rigidification-causing mutants display a clear trend of significantly decreased enzymatic activity and even complete loss in some cases (Fig 2). For example, G166A mutant retained some activity but G166P mutant (the most rigid mutation) almost completely lost activity. A similar situation is seen in 130’s loop. Single glycine to alanine mutants exhibited partial activity, while mutations of two glycines to alanines resulted in almost complete loss of activity, just like the R167A negative control. The results of huATCase mutants are consistent with the *E. coli* mutants except for a very small difference that a single mutation (G132A) can completely abolish activity (Fig 2C and F).

**Figure 2.**
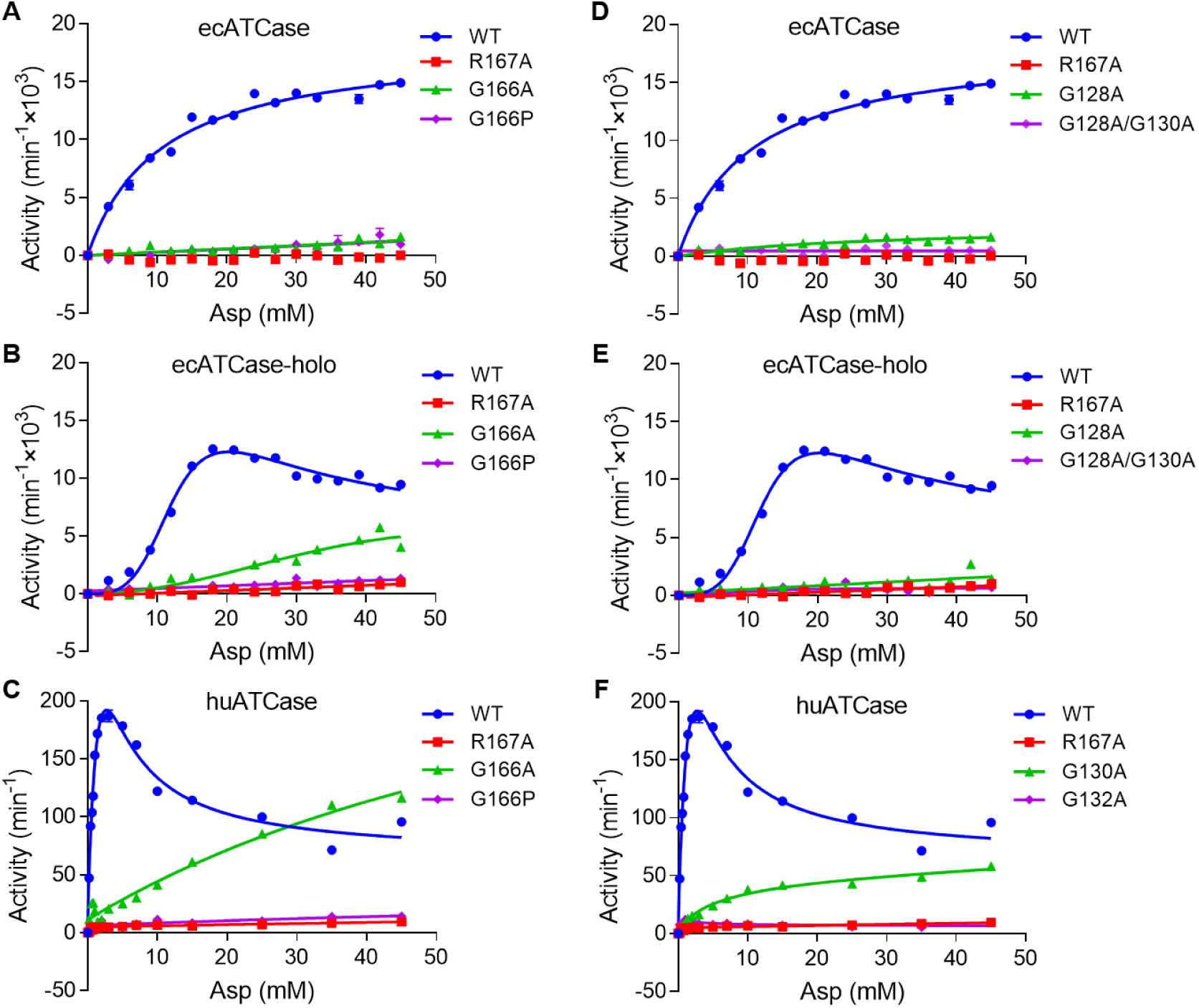
Enzyme kinetics curve of different mutants of ecATCase, ecATCase-holo, and huATCase. In each graph, corresponding wild-type and R167A ATCase were used as positive and negative control, respectively. ATCases used for each group are: ecATCase (**A**, **D**), ecATCase-holo (**B**, **E**) and huATCase (**C**, **F**).

To further confirm the importance of the flexibility of R167/130’s loop, we “locked” ecATCase and ecATCase-holo at R state by using the C47A/A241C mutants of ecATCase and ecATCase-holo as previously reported (Mendes & Kantrowitz, 2010a, Mendes & Kantrowitz, 2010b, West, Tsuruta et al., 2002). The enzymatic activity result is almost the same; G166P and G128A/G130A mutants lost almost all activity (Fig EV1). Taken together, we conclude that the flexibility of R167/130’s loop is important for ATCase’s catalytic function, including that at the R state.

### The flexibility of R167/130’s loop has a close relationship with K_m_ value of ATCase

Based on enzyme kinetics curves, V_max_, K_m_, and n_H_ were calculated and listed for various ATCases in Table EV1, which shows a strong correlation between K_m_ and the flexibility of R167/130’s loop. By analyzing the sequences and interactions of R167/130’s loop, we found that huATCase possesses the most flexible R167/130’s loop, owing to an additional glycine (G132) in 130’s loop (Fig 1D) and fewer interactions of R167 (Fig EV2 and Appendix Table S2). In comparison, ecATCase-holo possesses the least flexible R167/130’s loop, owing to more interactions of R167 and the additional interactions of 130’s loop derived from the hydrogen bond network at the interface between active site and the regulatory subunit.

The difference in flexibility is reflected in K_m_ values of various ATCases: huATCase has the smallest K_m_ value while ecATCase-holo has the largest. For ecATCase-holo locked in R state, K_m_ value dramatically decreased, even smaller than ecATCase, which indicates a more flexible R167/130’s loop. The K_m_ value did not change much after ecATCase was locked in R state, which is consistent with previous studies of ecATCase (Mendes & Kantrowitz, 2010a) and can be explained since ecATCase locked at R state cannot resemble a true R state ecATCase-holo due to the lack of regulatory subunits. Taken together, the flexibility of R167/130’s loop can notably influence catalytic property of ATCase in both human and *E. coli* enzymes, and ATCase with a more flexible R167/130’s loop, would be more sensitive to the change of substrate concentration and easier to achieve full catalytic activity.

### ATCase mutants with a rigid R167/130’s loop restrict R167 at either “out” or “in” state

To further study the flexibility of R167/130’s loop, we managed to solve the structures of G166P and G128A/G130A mutants of ecATCase and ecATCase-holo. Data collection and refinement statistics are shown in Table EV2. Corresponding mutations were confirmed in the electron density maps (Fig 1C). As shown in Fig 1C, R167 of G166P ecATCase and ecATCase-holo is restricted at “out” and “in” state, respectively. The situation is similar in the case of G128A/G130A ecATCase and ecATCase-holo. Given the fact that all these ATCase variants lost their activity almost completely, we conclude that neither R167 “in” nor R167 “out” state alone is sufficient for the catalytic function of ATCase and R167 needs to be able to switch between “in” and “out” state in the catalytic cycle. Additionally, due to the close proximity and multiple interactions between R167 and 130’s loop (Fig EV2), R167’s flexibility is largely restricted if 130’s loop is rigid, which explains why the flexibility of 130’s loop is important and necessary.

### ATCase mutants with rigid R167/130’s loop can bind CP but cannot further bind Asp

To further assess the significance of R167’s conformation switch between “in” and “out” state during ATCase catalytic process, we did ITC experiments using wild-type, R167A, G166P, and G128A/G130A mutants of the ecATCase, in which wild-type and R167A mutant were the positive and negative control, respectively. We tested the binding of the ATCase enzymes with the natural substrates, CP and Asp. Our results show that all ATCase variants were able to bind CP, meaning that these mutations do not affect CP binding (Fig EV3, top). After CP binding, we titrated Asp in ATCase. For wild-type ecATCase, the reaction heat was so large, indicating enzymatic reaction, and the binding heat was masked completely (Fig EV3A, bottom). For ATCase mutants, only very small heat peaks appeared (Fig EV3B-D, bottom), indicating no enzymatic reaction occurred, which is consistent with the results of enzymatic kinetics assays. In the meanwhile, heat peaks in each assay are of almost the same height, indicating no Asp binding occurred. We also performed ITC assays using ecATCase-holo and the results are the same with ecATCase (Fig EV4). All calculated ITC parameters are listed in Appendix Table S3. Because the mutants of ecATCase and ecATCase-holo have been shown to be either “locked” at R167 “in” or “out” state, it is clear that the flexibility afforded by R167/130’s loop is essential in helping Asp enter the active site to enable catalytic function.

### Molecular dynamics simulation of R167 switch from “in” to “out” state of ATCase

Next, we performed a molecular dynamic simulation, in which one catalytic chain was chosen for each energy calculation and MD simulation. First, we calculated the total energy of R167 “in” and “out” state of huATCase (PDB ID: 5G1N and 5G1O), ecATCase (PDB ID: 1EKX and 3CSU), and ecATCase-holo (PDB ID: 4KGV and the wild-type apo-ecATCase-holo structure solved in this paper), in which PALA binding form ATCase were used for R167 “in” state and apo form ATCase were used for R167 “out” state. It was found that the energy difference between the two states in huATCase is smaller than ecATCase or ecATCase-holo (Fig EV5), which suggests that R167 may be easier to switch in huATCase. This is consistent with our analysis demonstrating that huATCase possesses more flexible R167/130’s loop. We also calculated the energy of apo-ecATCase-holo (PDB ID: 4FYW) with R167 “in” state, and found it is close to and even higher than the energy of ecATCase-holo with R167 “out” state, indicating this structure may be an easier one to observe R167 switch in ecATCase-holo.

For MD simulation, the PALA bound structures (PDB ID: 5G1N, 1EKX, and 4KGV) with R167 “in” state were used firstly and PALA was removed in each model, which would facilitate “in” to “out” transition switch. For huATCase, after 20 ns simulation, R167 was able to switch from “in” to “out” state. During this simulation, huATCase domain opening took place, followed by gradual change of R167 from “in” to “out” state accompanied by the conformational change of 130’s loop (Movie EV1). The final conformation of 130’s loop was highly consistent with that in apo-huATCase (PDB ID: 5G1O). However, for ecATCase and ecATCase-holo, we did not observe this switch after 100 ns, which is consistent with the energy analysis above. We thus further performed the same simulation using apo-ecATCase-holo with R167 “in” state (PDB ID: 4FYW) and observed R167 switch after 40 ns (Movie EV2). The start and end models in simulations where R167 switch occurred were aligned and are shown in Fig 3A and B. The heat maps depicting the cross-correlation of the Cα of residues are shown in Fig EV5B.

**Figure 3.**
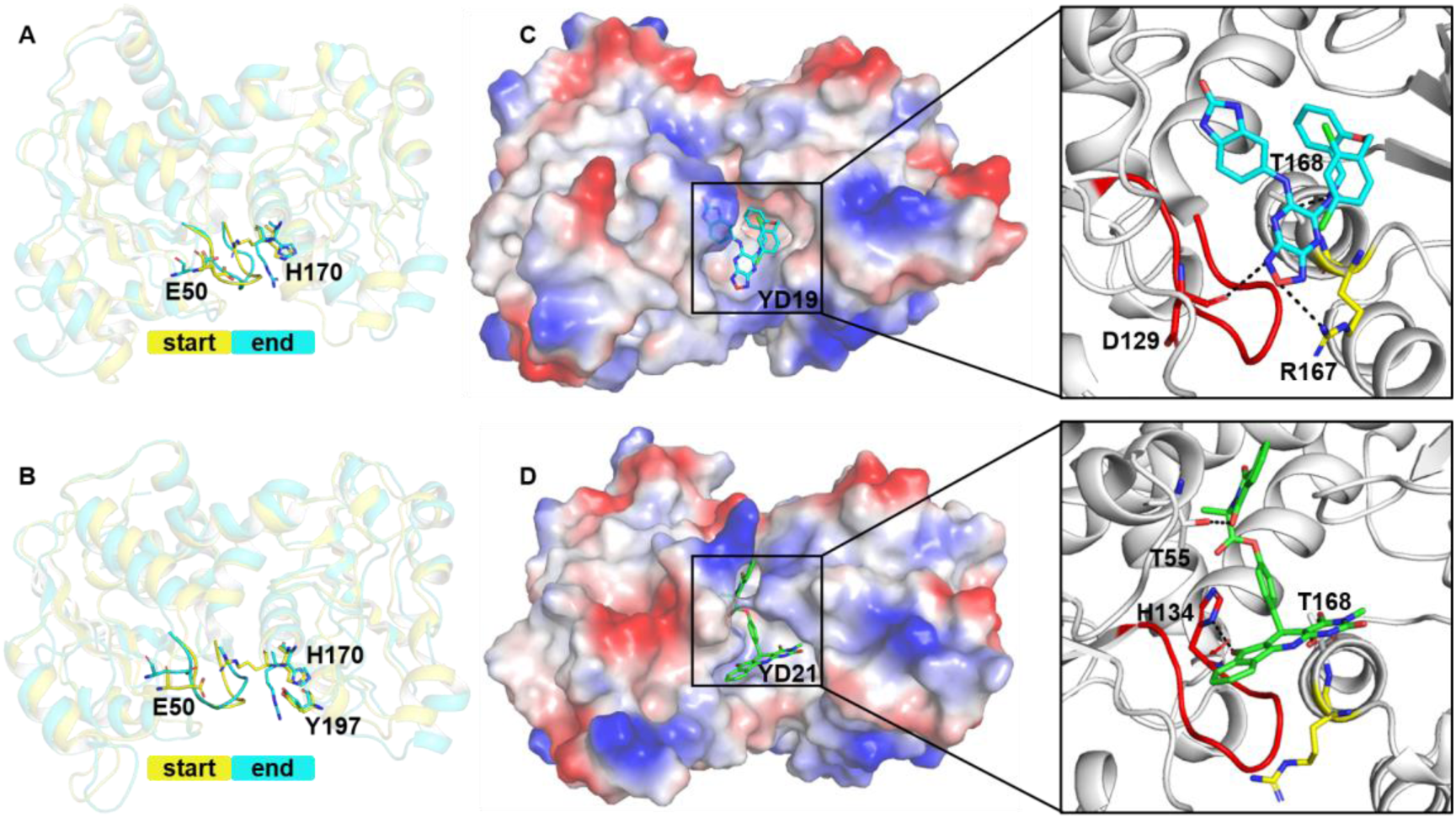
MD simulation of R167 switch from “in” to “out” state and binding models of YD19 and YD21 with huATCase. **A**, **B** Structural comparison of the start and end models of the MD simulation for R167 switch in huATCase (PDB ID: 5G1N) and ecATCase (PDB ID: 4FYW), respectively. Important residues interacting with R167 in the conformational switch are labeled and shown as sticks. The switch is shown visibly in Movie EV1 and Movie EV2. **C**, **D** The detailed binding models and interactions of YD19 and YD21 with huATCase. Compounds are shown as sticks together with transparent electrostatic surface of the protein (left). Residues involved in polar interactions with compounds are shown as sticks and labeled in black (right).

### The R167/130’s loop region is closely related to the feedback regulation of ATCase

We carried out a fluorescent assay to further demonstrate that ATCase possessing a rigid region of R167/130’s loop is not able to transit from T to R state. ecATCase-holo was used in this experiment and results are shown in Appendix Fig S2. Consistent with our ITC results, only wild-type ecATCase-holo was able to undergo T to R transition, whereas G166P and G128A/G130A mutants could not, akin to R167A mutant (Appendix Fig S2B). This result reveals that this region likely controls the difficulty level for ATCase to transit from T to R state, which is also regulated by the binding of different nucleotides in the feedback regulation. In light of the fact that this R167/130’s loop region locates at the interface between active site and regulatory subunit, we consider that it may serve as a previously unknown feedback regulatory feature in ecATCase-holo function.

To verify our speculation, we performed MD simulation using ecATCase-holo (one catalytic chain and one regulatory chain were used) to detect the structural difference around the R167/130’s loop region as a result of pyrimidines or purines binding. A previous structure (PDB ID: 4FYY) (Cockrell & Kantrowitz, 2012) was chosen for the pyrimidines binding model of T state ecATCase-holo; and purines binding model was obtained by replacing the pyrimidines by purines in the same structure. The pyrimidines and purines binding models of R state ecATCase-holo were also established based on the relevant structures (PDB ID: 4KH1 and 4KH0) (Cockrell et al., 2013). After 20 ns simulation, we found that for T state ATCase ecATCase-holo, the binding free energy of pyrimidines or purines binding model between catalytic and regulatory subunit displayed a significant difference. Comparing with pyrimidines binding model, purines binding caused a higher binding free energy, indicating a less stable combination between catalytic and regulatory subunit, and the hydrogen bond network associated with the region of R167/130’s loop was also partially destroyed, which was not found in R state ecATCase-holo (Appendix Fig S3). Taken together, these results suggest a close relationship between the region of R167/130’s loop and the feedback regulation.

### Virtual compound screening yields two inhibitors targeting apo-huATCase

Since huATCase is a known cancer drug target, we wondered whether the newly found R167/130’s loop region of ATCase could be targeted, in conjunction with the active site, to develop new dual-targeting inhibitors for ATCase. To this end, we performed a virtual compound screening simultaneously targeting both the active site and the newly found regulatory region of apo-huATCase. After two rounds of screening, 27 high-ranking compounds were selected and purchased in a small amount. We then performed 5 rounds of preliminary inhibition experiments for huATCase and selected 5 compounds (YD9, YD11, YD19, YD20, and YD21) which showed strong and consistent inhibition on the activity of ATCase. Further experiments helped us determine 2 decisions (YD19 and YD21) finally. The whole computer-aided screening workflow is shown in Fig 4A.

**Figure 4.**
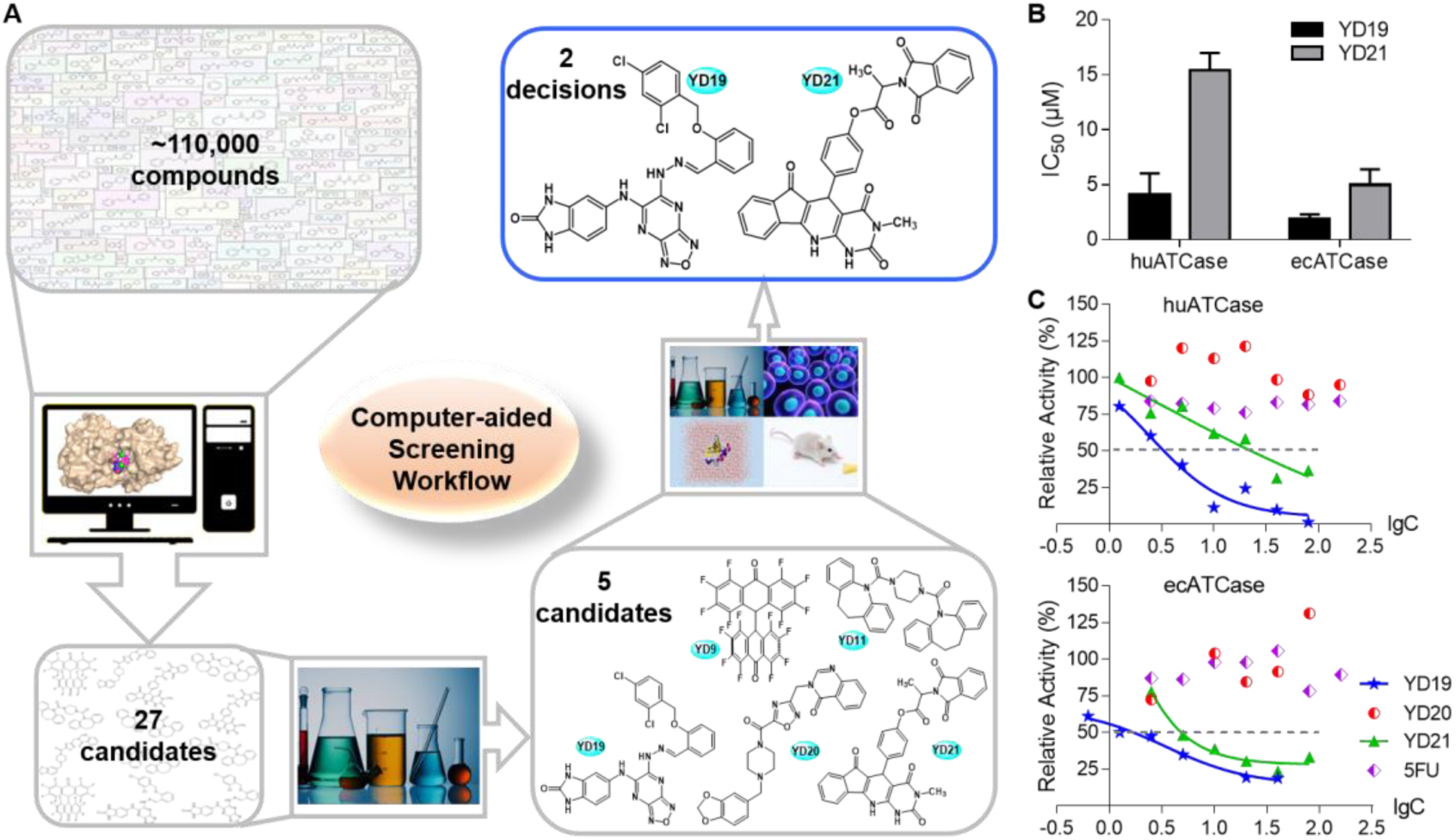
Virtual compound screening workflow and enzyme inhibition assays of YD19 and YD21 compounds. **A** Computer-aided screening workflow. The chemical structures of five candidates and two final decisions (in the blue rounded rectangle) are shown. **B** IC_50_ value of YD19 and YD21 for huATCase and ecATCase derived from **C**. **C** IC_50_ determination of YD19 and YD21 for huATCase and ecATCase. Datasets of YD19 and YD21 were fitted with Dose-response equation and inhibition at 50% is shown as a dashed line. YD20 was also tested and 5FU was used as a negative control in each graph.

After the 5 candidates were determined, we purchased a large quantity of these 5 compounds and carried out quantitative inhibition experiments. YD9 and YD11 were quickly abandoned due to their poor solubility, and YD19, YD20, and YD21 were used for the experiments. As shown in Fig 4B, YD19 and YD21 stood out with IC_50_ of 4.1 ± 1.9 μM and 15.4 ± 1.6 μM, respectively. We also tested the IC_50_ of these two compounds for ecATCase, which were 1.8 ± 0.4 μM and 5.0 ± 1.4 μM. YD20 and Fluorouracil (5FU) had no significant inhibition (Fig 4C); 5FU is a known cancer drug and will be used as the positive control in our MTT cell toxicity assays. ITC assays detecting the binding of these four compound with ATCase also produced consistent results, in which YD19 and YD21 showed binding to ecATCase and huATCase, whereas YD20 and 5FU did not (Appendix Fig S4). Calculated ITC parameters are listed in Appendix Table S3.

### Docking YD19 and YD21 to huATCase

After identifying YD19 and YD21 as top candidate inhibitors, we performed a more vigorous docking study. The two compounds can adopt 4 configurations due to tautomerism and cis-trans isomerism in YD19 and optical isomerism in YD21 (Appendix Fig S5A and C), respectively. Thus, we performed docking for all 4 configurations of each compound, followed by molecular simulation which was heated and equilibrated for 50 ns. According to the binding free energy analysis (Appendix Fig S5B and D), the best binding model of each compound and corresponding interactions are shown in Fig 3C and D. YD19 interacts with D129, R167 and T168 and YD21 interact with T55, H134 and T168. YD19 appears better than YD21 because it rigidifies the R167/130’s loop region by interacting with it and its binding is also more stable, according to the binding free energy results.

### YD19 and YD21 inhibit the proliferation of several cancer cell lines in MTT assay

To evaluate the anticancer potential, we performed cytotoxicity studies of the two compounds using six cell lines, including five cancer cell lines (A549, Hela, MCF7, HepG2, PC3) and one normal somatic cell line (CCC) using MTT assay, with 5FU as a positive control. As shown in Fig 5A, the cytotoxicity of the compounds varies in different cell lines. YD19 has good inhibitory effect on Hela, MCF7, HepG2, and PC3, whereas YD21 has an appreciable inhibitory effect on all six cell lines. In general, for cancer cell lines YD19 and YD21 are better than the clinically used anticancer drug 5FU, while YD19 is a slightly better than YD21 except for A549 cells; for normal cell lines (CCC), YD19 has the least toxicity. Therefore, YD19 seems a better molecule among the two candidate compounds and control. For comparison, YD20 was also tested at a single concentration but it could not effectively inhibit all six cell lines (Appendix Fig S6), which is consistent with its poor inhibition of ATCase catalytic activity.

**Figure 5.**
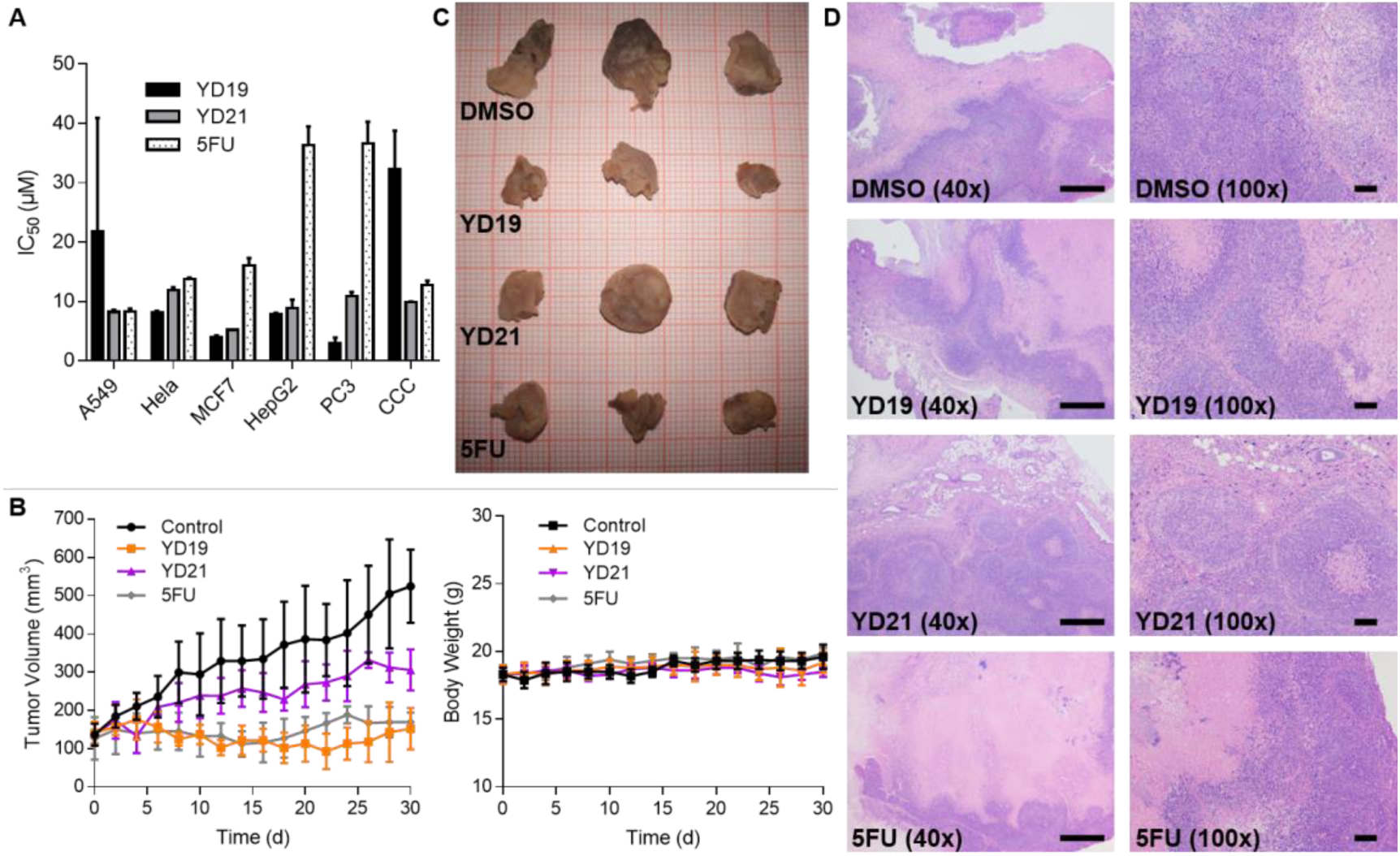
Results of MTT cytotoxicity assay and xenograft mouse assay. **A** MTT cytotoxicity result of YD19, YD21, and 5FU in six cell lines. See Appendix Fig S6 for full description of these cell lines. **B** Tumor volume (left) and body weight (right) change of mice in different groups via i.t. injection once every 2 days for total of 15 treatments. **C** Final tumor pictures of different groups. **D** Hematoxylin and eosin staining of tumor section in each group. Photographs at left and right were amplified 40× (with a ruler 500 μm) and100× (with a ruler 100 μm), respectively.

### YD19 and YD21 inhibit tumor growth in xenograft assays

BALB/c (nu/nu) mice with xenograft Hela tumor in the flanks were randomized into four groups and treated with DMSO, YD19, YD21, and 5FU respectively via i.t. injection every 2 days for a month. As shown in Fig 5B, YD19 and YD21 both inhibited the growth of xenograft tumors similar to 5FU; YD19 was more effective than YD21. The weights of mice were not affected by these compounds, which may be explained by the i.t. injection method we used. The final tumor volume in YD19 group was notably smaller than the DMSO group, and a similar situation occurred in 5FU group but not in YD21 group (Fig 5C). Hematoxylin and eosin staining of tumor sections showed extensive death of cancer cells in YD19, YD21, and 5FU groups. Cancer cells only occupied a small part of the whole tumor tissue and were restricted focally, indicating very weak diffusion. In contrast, in the negative control DMSO group, cancer cells occupied a larger portion of the entire tumor tissue and showed a dispersive distribution, indicating relative strong diffusion (Fig 5D). These results demonstrate that the two compounds are promising in not only impeding the growth and proliferation of multiple cancer cell lines *in vitro* but also inhibiting tumor growth *in vivo*.

## Discussion

In this work, motivated by our newly discovered feedback regulatory mechanism, we have successfully identified inhibitory compounds using a dual-targeting strategy. The lead compounds have demonstrated promise in enzymatic assay, in vitro, and in vivo. A model depicts the whole work is shown in Fig 6.

**Figure 6.**
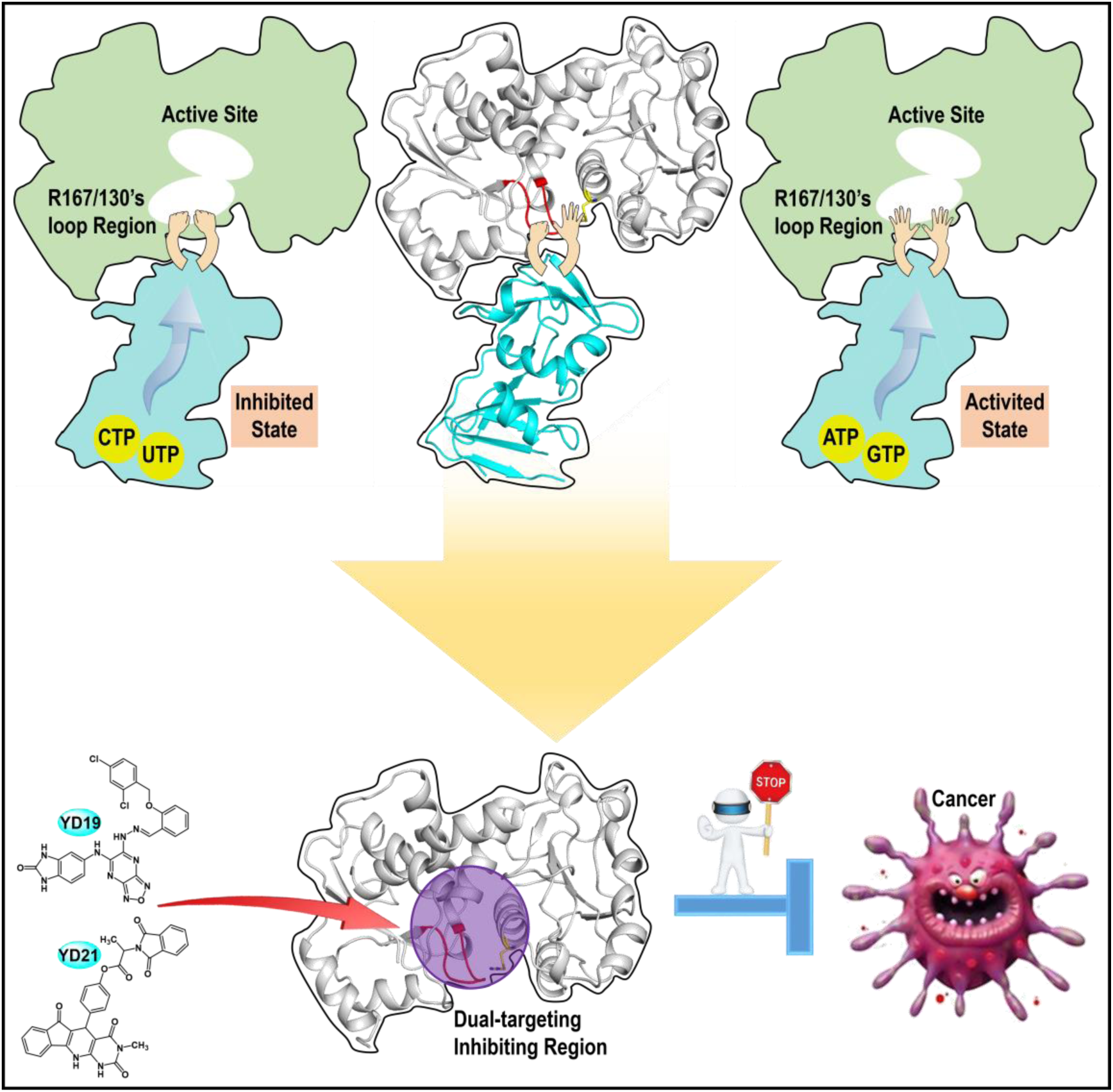
A model of newly discovered feedback regulatory mechanism of ATCase and the resulting dual-targeting strategy for developing potential anticancer drugs. The R167/130’s loop region located at the interface acts as a modulator between regulatory subunit and active site of ATCase, in response of the binding of pyrimidines or purines, which will further affect the active site, resulting in either inhibited or activated state of ATCase (top). Based on the newly found mechanism, a dual-targeting strategy was applied in developing potential anticancer drugs targeting huATCase, and the dual-targeting region was indicated by a semitransparent purple circle (bottom).

During the study on ATCase, we firstly solved a wild-type apo-ecATCase-holo with R167 “out” state (Fig 1A), which has helped uncover a previously neglected regulatory region of ATCase including R167 and 130’s loop. Through mutagenesis, we were able to reduce the conformational flexibility of R167/130’s loop and facilitate “out” state in ecATCase and “in” state in ecATCase-holo respectively (Fig 1C). Using both *E. coli* and human ATCase mutants as a probe, we revealed that neither R167 “in” nor “out” state alone is adequate to enable ATCase catalytic function as evidenced by our enzymatic assay and ITC assay results. During ATCase catalytic cycle, R167 needs to switch between “in” and “out” states, modulated by 130’s loop, which help Asp enter the active site of ATCase and very likely to help the release of product CA, too. 130’s loop is further modulated by regulatory subunit in the case of ecATCase-holo. Therefore, the flexibility of R167/130’s loop region plays a key regulatory role in the catalytic process of ATCase.

Our finding that there is a correlation between K_m_ value and flexibility of R167/130’s loop is very intriguing. K_m_ value is smaller for ATCase with more flexible region of R167/130’s loop, indicating it is more sensitive to the change of substrate concentration and easier to achieve full catalytic activity. MD simulating R167 switch from “in” to “out” state also shows consistent results. Another factor can notably influence the K_m_ value is the type of nucleotides, in which K_m_ value increases with pyrimidines bound and decreases with pyrimidines bound. Considering that the R167/130’s loop region is located between the active site and the regulatory subunit, we explored this region by MD simulation and found that there is a close relationship between the region and the feedback regulation. This conclusion is also supported by previous literature that mutating residues involved in the hydrogen bond network either destabilizes T state to promote R state of ecATCase-holo (K143rA mutant) (Eisenstein, Markby et al., 1990), or even abolishes the feedback effect of pyrimidines or purines (N111rA, N113rA and E142rA mutants) (Eisenstein et al., 1989).

Based on the findings mentioned above, we hypothesized the R167/130’s loop region as a previously unappreciated regulatory element in response to the binding of pyrimidines or purines, in which the binding of pyrimidines in regulatory subunit rigidifies this region while binding of purines relaxes it. Such changes in the region would further make T to R transition easier or more difficult, which represents the mechanism of the feedback regulation (Fig 6, top). In addition, we found the results of huATCase were very similar to ecATCase as evidenced by enzymatic assays and MD simulations; it is known that CAD is also regulated by cooperativity effect and feedback regulation (Moreno-Morcillo et al., 2017, Serre et al., 2004). Therefore, we inferred this mechanism in CAD, which laid foundation for us to design new inhibitors targeting apo-huATCase that would not cause domain closure as causing the failure of PALA. Building on the discovery of the new feedback regulation mechanism, we have successfully identified two inhibitors targeting both the newly found regulatory region and the active site of apo-huATCase (Fig 6, bottom). The compound position and extensive contacts with the R167/130’s loop region would make it almost impossible for R167 to switch from “out” to “in” state and interact with E50. Thus, after binding with these two inhibitors, domain closure of huATCase would not occur. The IC_50_ of the two compounds is micro-molarity (Fig 4B), which are significantly better than the existing inhibitors of apo-ecATCase (Heng, Stieglitz et al., 2006) (with a best IC_50_ of 79 μM, about 40-fold less potent than the best result we obtained). It is noted that owing to the relatively poor solubility and multiple configurations of the two compounds, the real inhibiting capacity of them may have been considerably stronger.

The two inhibitors derived from our dual-action strategy, which simultaneously target both the active site and the new feedback regulatory site of R167/130’s loop, represent a novel avenue to design anticancer drugs towards huATCase. Those initial compounds without any structural modification yet have already shown great promise as shown by our results of MTT and xenograft assays. They inhibit the proliferation of multiple cancer cell lines *in vitro*, as well as the growth of mice xenograft tumors *in vivo* (Fig 5). MD simulation and binding free energy analysis have helped us identify the best binding mode of each compound, which makes it possible to analyze the interactions. These results will certainly help guide chemical modifications of the compounds. Between the two lead compounds, YD19 is a better inhibitor and has better fit in the ATCase structure, thus representing a good starting point for structure modification. For clarity, we divide YD19 into three parts (Appendix Fig S7), in which part I occupies the active site region, part II occupies the newly found regulatory region and part III occupies the remaining region of the pocket. For part I, we would like to increase electronegativity to strengthen its interaction with the positive active site. While modification of part II can be minor, major modification can be applied in part III because the chlorophenyl moiety seems to be somewhat redundant. Other smaller substituent groups should be tested. Design and synthesis of new compounds are on the way.

## Materials and Methods

### Cloning, expression, and purification of ecATCase, ecATCase-holo, huATCase, and corresponding mutants

The cDNA of wild-type ecATCase and regulatory chain of ecATCase-holo were amplified by PCR (Qiagen Kit) using BL21(DE3) strain genome as template, and were inserted into pET28b and pET22b, respectively. The cDNA of wild-type huATCase was obtained as a gift from Han lab in Xiamen University, and was inserted into pOPINM (addGene) as reported by Ruiz-Ramos *et al*. (Ruiz-Ramos, Lallous et al., 2013). Site-directed mutation kit (Qiagen) was used to obtain plasmids with mutations using corresponding wild-type plasmids as templates. BL21(DE3) strain was chosen for expressing ecATCase and ecATCase-holo, and BL21(DE3)pLysS was used for expressing huATCase. Transformants were cultured in 1 L TB medium at 310 K and induced by 0.5 mM IPTG when OD_600_≈1.0, followed by overnight culturing at 289 K. Bacteria pellet was collected by centrifuging and resuspended in Buffer A (50 mM Tris-HCl pH 8.0, 300 mM NaCl and 10% Glycerol) for lysis by sonication. The lysate was then centrifuged at 15 000 ×g and the supernatant was added to the 1 mL Ni-NTA resin (Qiagen). After washing with Buffer A supplied with 30 mM imidazole, protein was eluted with 15 mL Buffer A supplied with 300 mM imidazole. The eluted protein was then buffer exchanged into Buffer B (50 mM Tris-acetate pH 8.3) for enzymatic activity and ITC assays, or Buffer C (50 mM Tris-acetate pH 8.3, 2 mM DTT and 5% Glycerol) for subsequent purification by HiLoad Superdex 200 column (GE). Protein in peak fractions was collected for crystallization assays.

### Crystallization and structure determination of ecATCase and ecATCase-holo

The preliminary crystallization condition was screened by the sparse matrix method and hanging drop vapor diffusion method was then used to improve the quality of preliminary crystal hits. The final optimal crystallization condition was 0.2 M NH_4_Ac, 0.1 M Tris pH 8.5, 20% PEG3350, and 10% glycerol for ecATCase, and 0.1 M HEPES pH 7.0, 30% Jeffamine M-600 pH 7.0, and 10% glycerol for ecATCase-holo. Crystals appeared in two days and grew to full size within ten days. X-ray diffraction data were collected using BL17U1 Beamline of Shanghai Synchrotron Radiation Facility (Wang, Zhang et al., 2018) at 0.979 Å or Rigaku X-ray generator at 1.542 Å. Datasets were processed by HKL-2000 (Otwinowski & Minor, 1997) and molecular replacement was performed by using a previous T state ecATCase-holo structure (PDB ID: 1ZA1) (Wang, Stieglitz et al., 2005) as searching template. Refinements were carried out by phenix.refine within Phenix (Adams, Afonine et al., 2010) and refmac5 within CCP4 suite (Collaborative Computational Project, 1994), as well as Coot (Emsley & Cowtan, 2004) for manual adjustments.

### Enzymatic activity assay of ATCase

Enzymatic activity assay was performed colorimetrically as previously reported (Pastra-Landis, Foote et al., 1981) and protein concentration was adjusted to make the final readout fall into rational range, which is 6 nM for ecATCase and ecATCase-holo, and 600 nM for huATCase. Final readout was determined by a microplate reader (Thermo) in 96-well plates and data were transformed into product concentration according to the standard curve, derived from the same approach using N-carbamoyl-DL-aspartate (TCI) as a standard reaction product (Appendix Fig S8). Datasets were fitted with the Michaells-Menten equation with/without substrate inhibition modification or the Hill equation with/without substrate inhibition modification as previously reported (Pastra-Landis, Evans et al., 1978), according to different situations. To calculate V_max_, K_m_, and n_H_, data at high concentration of substrate were truncated to eliminate the effect of substrate inhibition and fitted with Michaells-Menten or Hill equation. Paremeters and corresponding standard errors were calculated from these equations by OriginPro 2018 (Table EV1) and figures were plotted by GraphPad Prism 7.00. The concentration of different protein samples was measured by NonoPhotometer P-Class (IMPLEN) using their corresponding molar extinction coefficient (ε), in which the ε of ecATCase and ecATCase-holo were previously reported (Gerhart & Holoubek, 1967) and the ε of huATCase was calculated using ExPASy.

### Isothermal titration calorimetry

ITC assays for substrates binding were performed as follows. First, protein, Asp and CP were diluted to 50 μM, 500 μM, and 500 μM with Buffer B, respectively. For each variant of ecATCase and ecATCase-holo, three assays were done: 50 μM protein was titrated by 500 μM CP; 50 μM protein was titrated by 500 μM Asp; and 50 μM protein mixed with 4.8 mM CP was titrated by 500 μM Asp mixed with 4.8 mM CP. Data were processed by OriginPro 2018 to obtain parameters depicting the binding between substrates and ecATCase or ecATCase-holo.

ITC assays for inhibitors binding were performed as follows. First, different compounds (YD19, YD20, YD21, and 5FU) dissolved in DMSO were diluted to 500 μM with Buffer B, and final DMSO percentage was accurately controlled at 5%. Next, ecATCase and huATCase were diluted to 50 μM with Buffer B, in which process, 5% DMSO was added to ensure consistency with inhibitors. For both ecATCase and huATCase, four assays were performed that protein was titrated by YD19, YD20, YD21, and 5FU, respectively. Data were also processed by OriginPro 2018.

### Fluorescence assay

Fluorescence assays were performed as previously reported (Fetler, Tauc et al., 2001) with some modifications. Firstly, the two intrinsic tryptophan residues of ecATCase-holo were mutated to nonfluorescent phenylalanines. Next, rF145 (r indicates a residue in the regulatory chain of ecATCase-holo) was mutated to tryptophan to enable fluorescence signal during T to R transition. Enzymatic activity of G166P and G128A/G130A mutants based on W209F/W284F/rF145W were also tested to confirm consistency with preceding results (Appendix Fig S2A).

To detect fluorescence change during the T to R state transition of ecATCase-holo, following steps were performed. Protein (saturated with 4.8 mM CP) was loaded in a fluorescent cuvette and the excitation/emission wavelength was optimized. The final optimized wavelengths were 273 nm for excitation and 324 nm for emission, which were used for all time-course fluorescent assays. During these assays, the sample containing protein and CP was excited at 273 nm and the emission at 324 nm was continuously recorded for ∼20 s before a rapid injection of 30 mM Asp (final concentration), followed by a record for another ∼40 s. Final fluorescence signal change was obtained by substrating the signal in the blank control group from the sample groups.

### Virtual inhibitor screening

We performed virtual compound screening, targeting apo-huATCase, using AutoDock Vina (Trott & Olson, 2010) and AutoDockTools4 (Morris, Huey et al., 2009). A library containing ∼110,000 compounds (Pharmacodia Inc. Beijing) was obtained and those with the molecular weight (MW) greater than 1,000 were omitted. Search space was set at 30 Å × 30 Å × 30 Å, covering both the active site region and the newly identified R167/130’s loop region. Two rounds of screening were performed as follows. In the first round, no residue side chain of the receptor was treated as flexible during docking. Screening result was sorted by the docking score and the top 1,000 were selected for the second round. In the second round, residue side chains of receptor close to the docking compounds were treated as flexible and screening result was sorted by score. Next, compounds appearing in both the top 100 of the two rounds were compared and redundant structures were abandoned. Finally, the remaining compounds were purchased in a small amount for the inhibition assays.

### Enzymatic activity inhibiting assay of ATCase

For inhibition assays, substrate concentration at the V_max_ of the corresponding enzymatic kinetics curve was chosen, which is 30 mM Asp for ecATCase and 3 mM Asp for huATCase. Procedures are similar to the enzymatic activity assay except that different compounds were added before initiating the reaction with 4.8 mM CP. Experiment with the same percentage of DMSO was used as a control and all experiments also had a blank control without Asp to eliminate the additional absorption caused by different compounds.

For IC_50_ determination, compounds with relatively large quantity were needed and purchased (ChemDiv, California). For each compound, we carried out at least eight experiments using different concentrations in consecutive double dilution. Logarithms of compound concentrations were used as X value and datasets were fitted with dose-response equation. Corresponding IC_50_, as well as standard error, were calculated from the fitted equations by OriginPro 2018 and figures were plotted by GraphPad Prism 7.00.

### Molecular dynamics simulations

All MD simulations and post processes were performed using programs in Amber16 or AmberTools16 (Case, Betz et al., 2016). The same simulation protocol was used as follows. Firstly, tleap was used to generate the topology and coordinate files for each system, during which ff14SB force field parameters were used for protein, while parameters for small compounds were generated by antechamber and parmchk. Each system was neutralized by Na^+^ or Cl^−^ ions and was explicitly solvated by using the TIP3P water potential inside a box of water molecules with a minimum solute-wall distance of 10 Å, except for total energy calculation of a system, for which implicit solvated model was used instead of an explicit one. Next, pmemd was used to perform six cycles of minimizations to remove unfavorable contacts of each system, during which Cartesian restraints (decreasing from 0.1 kcal/mol/Å^2^ to 0) was applied to protein. The energy-minimized system was then heated over 200 ps from 0 to 310 K without restraints, during which constant volume was maintained. Finally, 2 ns unrestrained equilibration was carried out under constant pressure (1 bar) and temperature (310 K), followed by a 20-100 ns unrestrained molecular dynamics simulation. For post processes, Cpptraj was used to generate dynamic cross-correlation matrix and convert each frame of MD simulation into PDB format. MMPBSA.py was used to perform the binding free energy analysis, as well as the energy decomposition analysis.

### MTT cytotoxicity assay

All cell lines used in this research were obtained from the Cell Resource Center (Peking Union Medical College Headquarters of National Infrastructure of Cell Line Resource, NSTT). MTT assays were performed as follows. First, different types of cells were seeded into 96-well plates (1,000 cells/well) and cultured for 24 h. After adding compound, cells were continuously cultured for 3 d. Next, MTT solution was added and incubated in the dark for 4 h followed by careful removal of medium and addition of 150 μL DMSO. After shaking on a microplate reader for 10 min to adequately dissolve the Formazan reduced from MTT, readings at A570 nm was recorded and IC_50_ was calculated the same as referred above.

### Xenograft mouse model

The female BALB/c (nu/nu) mice were purchased from Vital River Laboratories (Beijing, China). All animal experiments were performed in accordance with the Guide for the Care and Used of Laboratory Animals and were approved by the Experimental Animal Ethics Committee in Beijing. For xenograft mouse assay, 5 × 10^6^ Hela cells were injected subcutaneously in the flanks of 20 four- to six-week-old female BALB/c (nu/nu) mice. After most of the tumor volumes exceeded 100 mm^3^, 12 mice with similar tumor volume were selected and randomly divided into four groups (3/group) with the treatment of 2.5 mg/kg DMSO (a negative control), YD19, YD21, and 5FU (a known cancer drug as a positive control) respectively via i.t. injection once every 2 days, lasting for one month. Tumor volume and body weight were measured every 2 days before injection. After 15 treatments, mice were euthanized, and the tumors were harvested, photographed, spliced, and stained by hematoxylin and eosin. The stained tumor splices were photographed and analysed under a microscope with a camera.

## Acknowledgments

We thank Han lab in Xiamen University for the generous gift of CAD cDNA and staff at BL17U1 beamline of Shanghai synchrotron facility for their help in diffraction data collection.

## Author contributions

Lei, Z. performed the main experiments and molecular dynamic simulations. Wang, N. contributed to X-ray data collection and structure determination. Wang, B. helped in mouse experiments. Lu, Z. helped in protein preparation. Tan, H. helped in dynamic simulations. Lei, Z., Wang, N., Zheng, J., and Jia, Z. designed the project and wrote the article. All authors reviewed and approved this article.

## Expanded View Figures, Tables and Movies

**Figure EV1.**
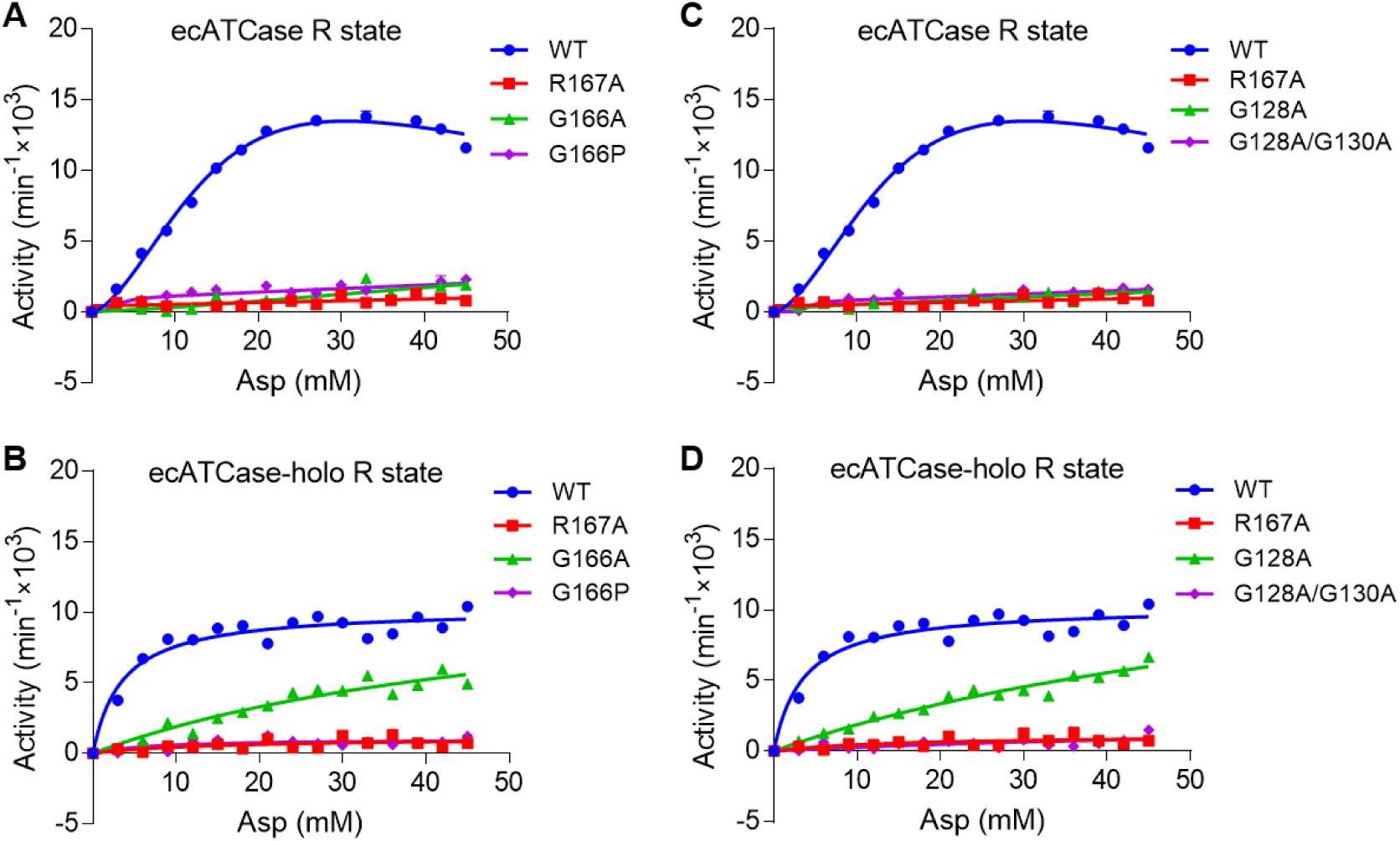
Enzyme kinetics curve of ecATCase or ecATCase-holo and their mutants locked at R state. In each graph, corresponding wild-type and R167A ATCase are used as positive and negative control, respectively. ATCases used for each group are: ecATCase locked at R state by C47A/A241C mutations (**A**, **C**) and ecATCase-holo locked at R state by C47A/A241C mutations (**B**, **D**).

**Figure EV2.**
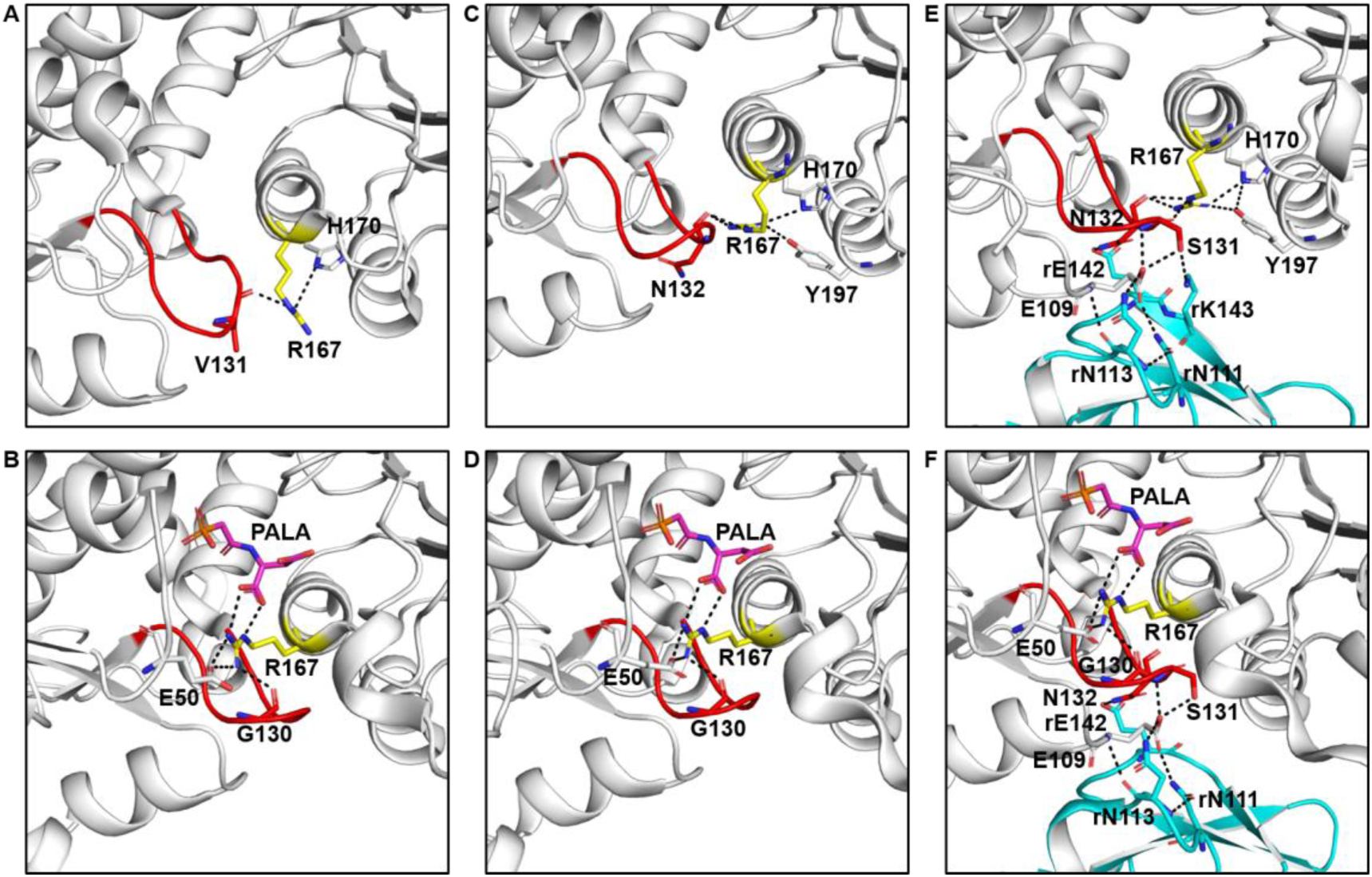
Important interactions with R167 and 130’s loop in various ATCases. In each graph, PALA (colored in magenta) or residues directly involved in the interactions are shown as stick and labeled in black. All interactions were listed in Appendix Table S2 ATCases used for each graph are: apo-huATCase (PDB ID: 5G1O, **A**), PALA-huATCase (PDB ID: 5G1N, **B**), apo-ecATCase (PDB ID: 3CSU, **C**), PALA-ecATCase (PDB ID: 1EKX, **D**), apo-ecATCase-holo solved in this work (**E**) and PALA-ecATCase-holo (PDB ID: 4KGV, **F**).

**Figure EV3.**
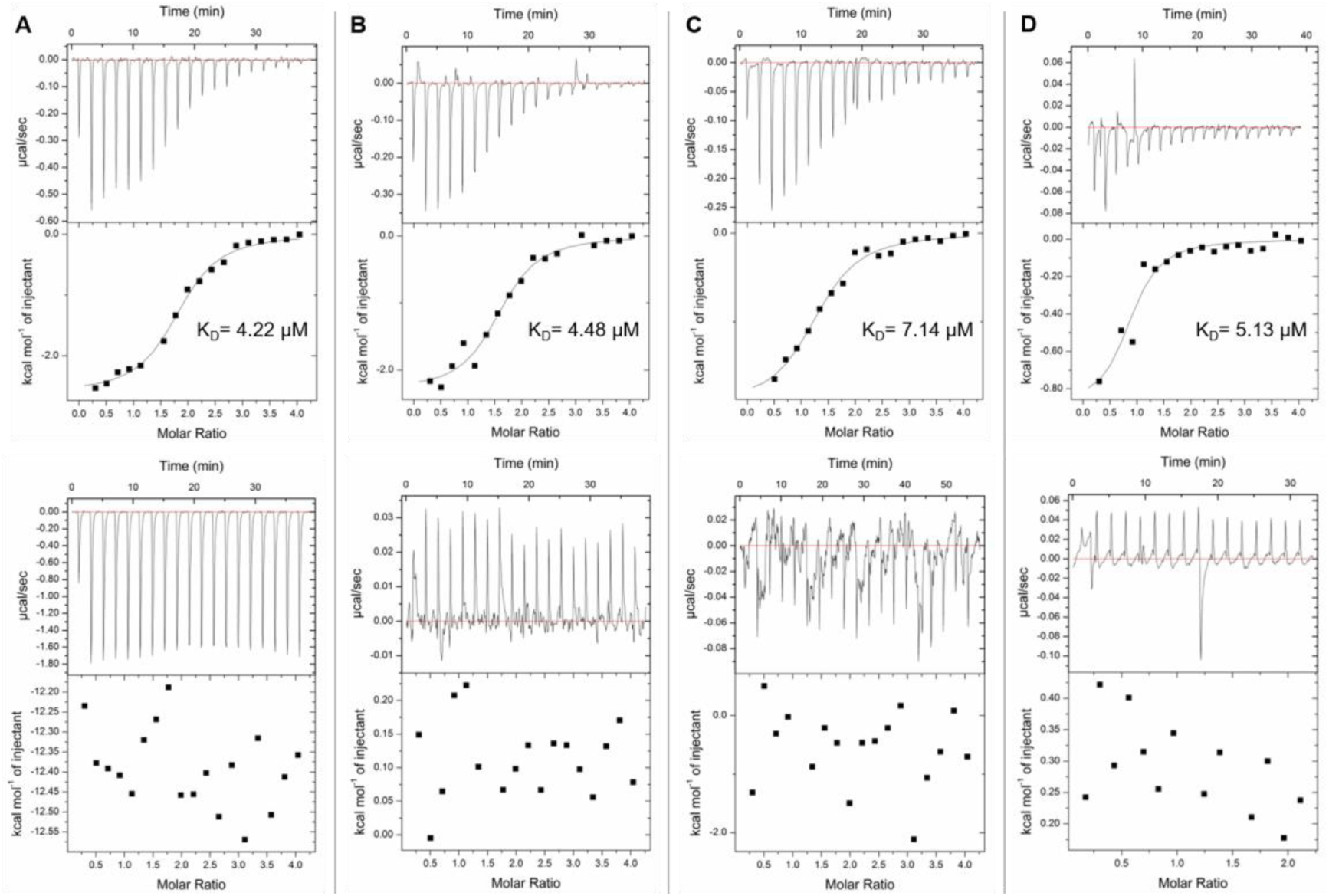
ITC results of ecATCase variants titrated by CP (top) and Asp after CP binding (bottom). In each assay, the concentration of CP and Asp used for titration is 500 μM, and ATCase is 50 μM. CP used to saturate ATCase is 4.8 mM. K_D_ is shown if binding curve can be fitted and other parameters were listed in Appendix Table S3. ATCases used for each group are: wild-type ecATCase (**A**), R167A ecATCase (**B**), G166P ecATCase (**C**) and G128A/G130A ecATCase (**D**).

**Figure EV4.**
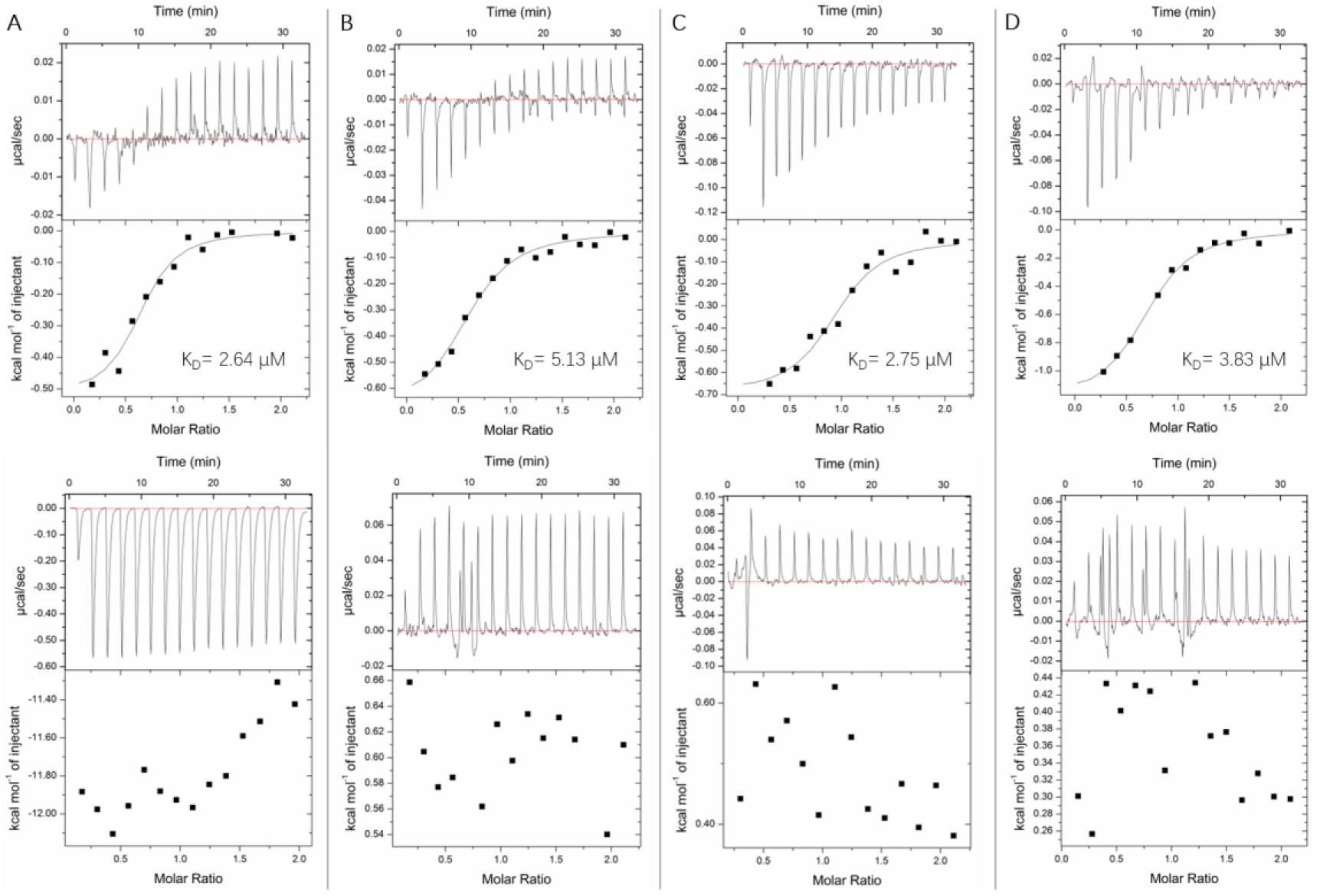
ITC results of ecATCase-holo variants titrated by CP (top) and Asp after CP binding (bottom). In each assay, the concentration of CP and Asp used for titration is 500 μM, and ATCase is 50 μM. CP used to saturate ATCase is 4.8 mM. K_D_ is shown if binding curve can be fitted and other parameters were listed in Appendix Table S3. ATCases used for each group are: wild-type ecATCase-holo (**A**), R167A ecATCase-holo (**B**), G166P ecATCase-holo (**C**) and G128A/G130A ecATCase-holo (**D**).

**Figure EV5.**
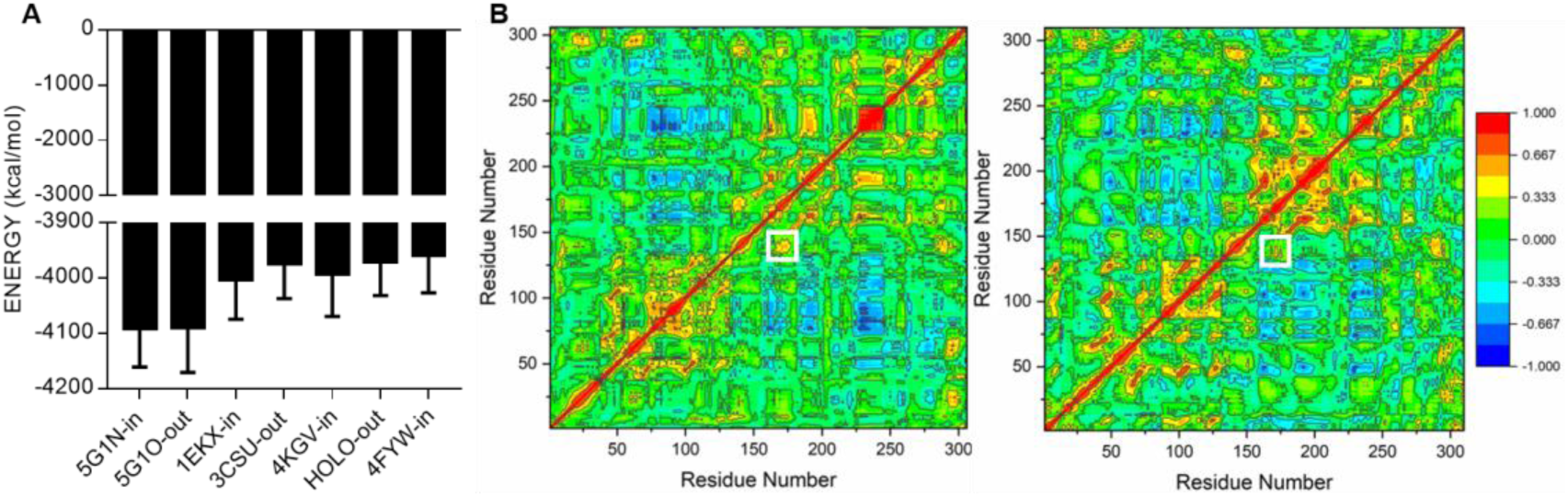
MD simulation of R167 switch from “in” to “out” state in huATCase and ecATCase. **A** Energy comparison of seven ATCases with R167 “in” or “out” state. The first two are huATCase, the middle two are ecATCase, and the last three are ecATCase-holo, in which the one named “HOLO-out” used the structure of wild-type apo-ecATCase-holo with R167 “out” state solved in this research and the last one used the wild-type apo-ecATCase-holo (PDB ID: 4FYW) with R167 “in” state. **B** Dynamic cross correlation heat map for R167 switch in huATCase (left, PDB ID: 5G1N) and ecATCase-holo (right, PDB ID: 4FYW). The white boxes indicate Cα correlation between R167 and 130’s loop.

**Table EV1.** V_max_, K_m_, and n_H_ of various ATCases.

| ATCase type | $V_{\max}$ ($\text{min}^{-1}$) | $K_m$ (mM) | $n_H^a$ |
| --- | --- | --- | --- |
| wild-type huATCase | $219.7 \pm 11.2$ | $0.4 \pm 0.1$ | 1 |
| wild-type ecATCase | $(18.3 \pm 0.7) \times 10^3$ | $10.0 \pm 1.6$ | 1 |
| wild-type ecATCase R state | $(14.6 \pm 0.5) \times 10^3$ | $10.2 \pm 0.6$ | $2.5 \pm 0.4$ |
| wild-type ecATCase-holo | $(16.1 \pm 3.3) \times 10^3$ | $12.6 \pm 1.4$ | $4.0 \pm 1.3$ |
| wild-type ecATCase-holo R state | $(11.5 \pm 0.8) \times 10^3$ | $4.9 \pm 1.1$ | 1 |

<sup>a</sup> $n_H = 1$ means this data set was fitted with Michaelis-Menten equation, while others were fitted with Hill equation.

**Table EV2.** Data collection and refinement statistics of five datasets of ecATCase or ecATCase-holo^a^.

| item | wild-type<br>holo | G166P | G166P<br>holo | G128A/G130A | G128A/G130A<br>holo |
| --- | --- | --- | --- | --- | --- |
| **Data collection statistics** |  |  |  |  |  |
| Wavelength<br>(Å) | 0.979 | 1.542 | 1.542 | 1.542 | 1.542 |
| Space group | R32 | R3 | P3<sub>2</sub>21 | P2<sub>1</sub> | P3<sub>2</sub>21 |
| Resolution<br>(Å) | 30.8-2.1<br>(2.1-2.1) | 36.5-2.8<br>(2.9-2.8) | 47.9-3.0<br>(3.1-3.0) | 47.1-2.5<br>(2.6-2.5) | 42.3-3.0<br>(3.1-3.0) |
| Unit cell |  |  |  |  |  |
| a, b, c (Å) | 129.7,<br>129.7,<br>198.0 | 128.9,<br>128.9,<br>48.3 | 126.6,<br>126.6,<br>196.5 | 81.8,<br>96.7,<br>121.7 | 127.4,<br>127.4,<br>197.5 |
| $\alpha, \beta, \gamma$ (°) | 90,<br>90,<br>120 | 90,<br>90,<br>120 | 90,<br>90,<br>120 | 90,<br>94,<br>90 | 90,<br>90,<br>120 |
| R<sub>meas</sub> | 0.127<br>(0.865) | 0.096<br>(0.768) | 0.180<br>(0.923) | 0.105<br>(0.702) | 0.113<br>(0.658) |
| Average (I/σ) | 15.8 (3.0) | 19.1 (2.5) | 13.1 (2.5) | 15.8 (2.6) | 12.8 (2.4) |
| Redundancy | 6.8 (7.3) | 3.3 (3.0) | 8.8 (8.6) | 4.1 (4.0) | 4.8 (4.5) |
| Completeness<br>(%) | 98.8 (95.8) | 91.6 (63.0) | 99.4 (97.2) | 99.2 (93.7) | 98.4 (95.8) |
| **Refinement statistics** |  |  |  |  |  |
| Resolution<br>(Å) | 30.8-2.1 | 36.5-2.8 | 47.9-3.0 | 47.1-2.5 | 42.3-3.0 |
| Reflections | 270867<br>(29096) | 22978<br>(2075) | 325738<br>(31375) | 270730<br>(25920) | 175524<br>(15995) |
| R<sub>work</sub>/R<sub>free</sub> | 0.18/0.21 | 0.21/0.26 | 0.21/0.24 | 0.19/0.23 | 0.24/0.27 |
| Number of atoms |  |  |  |  |  |
| protein | 3387 | 2202 | 10022 | 13601 | 10005 |
| zinc | 1 | 0 | 3 | 0 | 3 |
| water | 326 | 25 | 28 | 509 | 17 |
| RMS deviations |  |  |  |  |  |
| bond<br>lengths (Å) | 0.006 | 0.008 | 0.004 | 0.008 | 0.004 |
| angles (°) | 1.07 | 1.30 | 1.07 | 1.23 | 1.02 |
| Mean B value<br>(Å<sup>2</sup>) | 47.3 | 49.0 | 53.1 | 45.0 | 72.0 |

<sup>a</sup>Values in parentheses correspond to the highest-resolution shell.

**Movie EV1. MD simulation of R167 switch from “in” to “out” state in huATCase.**

In this movie, R167, E50, and H170 are shown as sticks, in which E50 and H170 interact with R167 at “in” and “out” state, respectively. R167 and 130’s loop were colored in yellow and red, respectively. During this simulation, it can be observed that domain opening took place first, followed by gradual change of R167 from “in” to “out” state accompanied by the conformational change of 130’s loop.

**Movie EV2. MD simulation of R167 switch from “in” to “out” state in apo-ecATCase-holo.**

In this movie, R167, E50, H170, and Y197 are shown as sticks, in which E50 and H170/Y197 interact with R167 at “in” and “out” state, respectively. R167 and 130’s loop were colored in yellow and red, respectively. During this simulation, it can be observed that R167 gradually switches from “in” to “out” state accompanied by the conformational change of 130’s loop.

## References

Adams PD, Afonine PV, Bunkoczi G, Chen VB, Davis IW, Echols N, Headd JJ, Hung LW, Kapral GJ, Grosse-Kunstleve RW, McCoy AJ, Moriarty NW, Oeffner R, Read RJ, Richardson DC, Richardson JS, Terwilliger TC, Zwart PH (2010) PHENIX: a comprehensive Python-based system for macromolecular structure solution. Acta crystallographica Section D, Biological crystallography 66: 213–21

Aoki T, Weber G (1981) Carbamoyl phosphate synthetase (glutamine-hydrolyzing): Increased activity in cancer cells. Science (New York, NY) 212: 463–464

Case D, Betz R, Cerutti DS, Cheatham T, Darden T, Duke R, Giese TJ, Gohlke H, Götz A, Homeyer N, Izadi S, Janowski P, Kaus J, Kovalenko A, Lee T-S, LeGrand S, Li P, Lin C, Luchko T, Kollman PA (2016) Amber 2016, University of California, San Francisco.

Cockrell GM, Kantrowitz ER (2012) Metal ion involvement in the allosteric mechanism of Escherichia coli aspartate transcarbamoylase. Biochemistry 51: 7128–37

Cockrell GM, Zheng Y, Guo W, Peterson AW, Truong JK, Kantrowitz ER (2013) New paradigm for allosteric regulation of Escherichia coli aspartate transcarbamoylase. Biochemistry 52: 8036–8047

Collaborative Computational Project N (1994) The CCP4 suite: programs for protein crystallography. Acta crystallographica Section D, Biological crystallography 50: 760–3

Eisenstein E, Markby DW, Schachman HK (1989) Changes in stability and allosteric properties of aspartate transcarbamoylase resulting from amino acid substitutions in the zinc-binding domain of the regulatory chains. Proceedings of the National Academy of Sciences of the United States of America 86: 3094–8

Eisenstein E, Markby DW, Schachman HK (1990) Heterotropic effectors promote a global conformational change in aspartate transcarbamoylase. Biochemistry 29: 3724–31

Emsley P, Cowtan K (2004) Coot: model-building tools for molecular graphics. Acta crystallographica Section D, Biological crystallography 60: 2126–32

Evans DR, Guy HI (2004) Mammalian pyrimidine biosynthesis: fresh insights into an ancient pathway. The Journal of biological chemistry 279: 33035–8

Fetler L, Tauc P, Herve G, Cunin R, Brochon JC (2001) Tryptophan residues at subunit interfaces used as fluorescence probes to investigate homotropic and heterotropic regulation of aspartate transcarbamylase. Biochemistry 40: 8773–82

Gerhart JC, Holoubek H (1967) The purification of aspartate transcarbamylase of Escherichia coli and separation of its protein subunits. The Journal of biological chemistry 242: 2886–92

Gerhart JC, Pardee AB (1962) The enzymology of control by feedback inhibition. The Journal of biological chemistry 237: 891–6

Gouaux JE, Lipscomb WN (1990) Crystal structures of phosphonoacetamide ligated T and phosphonoacetamide and malonate ligated R states of aspartate carbamoyltransferase at 2.8-. ANG. resolution and neutral pH. Biochemistry 29: 389–402

Gouaux JE, Stevens RC, Lipscomb WN (1990) Crystal structures of aspartate carbamoyltransferase ligated with phosphonoacetamide, malonate, and CTP or ATP at 2.8-. ANG. resolution and neutral pH. Biochemistry 29: 7702–7715

Grem JL, King SA, O’Dwyer PJ, Leyland-Jones B (1988) Biochemistry and clinical activity of N-(phosphonacetyl)-L-aspartate: a review. Cancer research 48: 4441–4454

Guo W, West JM, Dutton AS, Tsuruta H, Kantrowitz ER (2012) Trapping and structure determination of an intermediate in the allosteric transition of aspartate transcarbamoylase. Proceedings of the National Academy of Sciences of the United States of America 109: 7741–6

Ha Y, Allewell NM (1998) Intersubunit hydrogen bond acts as a global molecular switch in Escherichia coli aspartate transcarbamoylase. Proteins 33: 430–43

Heng S, Stieglitz KA, Eldo J, Xia J, Cardia JP, Kantrowitz ER (2006) T-state inhibitors of E. coli aspartate transcarbamoylase that prevent the allosteric transition. Biochemistry 45: 10062–71

Howlett GJ, Schachman HK (1977) Allosteric regulation of aspartate transcarbamoylase. Changes in the sedimentation coefficient promoted by the bisubstrate analog N-(phosphonacetyl)-L-aspartate. Biochemistry 16: 5077–5083

Huang J, Lipscomb WN (2004) Products in the T-state of aspartate transcarbamylase: crystal structure of the phosphate and N-carbamyl-L-aspartate ligated enzyme. Biochemistry 43: 6422–6

Huang J, Lipscomb WN (2006) T-state active site of aspartate transcarbamylase: crystal structure of the carbamyl phosphate and L-alanosine ligated enzyme. Biochemistry 45: 346–52

Jones ME (1980) Pyrimidine nucleotide biosynthesis in animals: genes, enzymes, and regulation of UMP biosynthesis. Annu Rev Biochem 49: 253–79

Kantrowitz E, Lipscomb W (1988) Escherichia coli aspartate transcarbamylase: the relation between structure and function. Science (New York, NY) 241: 669–674

Kantrowitz ER (2012) Allostery and cooperativity in Escherichia coli aspartate transcarbamoylase. Archives of biochemistry and biophysics 519: 81–90

Ke HM, Lipscomb WN, Cho YJ, Honzatko RB (1988) Complex of N-phosphonacetyl-L-aspartate with aspartate carbamoyltransferase. X-ray refinement, analysis of conformational changes and catalytic and allosteric mechanisms. Journal of molecular biology 204: 725–47

Krause KL, Volz KW, Lipscomb WN (1987) 2.5 A structure of aspartate carbamoyltransferase complexed with the bisubstrate analog N-(phosphonacetyl)-L-aspartate. Journal of molecular biology 193: 527–53

Ladjimi MM, Kantrowitz ER (1988) A possible model for the concerted allosteric transition in Escherichia coli aspartate transcarbamylase as deduced from site-directed mutagenesis studies. Biochemistry 27: 276–83

Lee L, Kelly RE, Pastra-Landis SC, Evans DR (1985) Oligomeric structure of the multifunctional protein CAD that initiates pyrimidine biosynthesis in mammalian cells. Proceedings of the National Academy of Sciences 82: 6802–6806

Lipscomb WN, Kantrowitz ER (2012) Structure and mechanisms of Escherichia coli aspartate transcarbamoylase. Accounts of chemical research 45: 444–53

Mendes KR, Kantrowitz ER (2010a) A cooperative Escherichia coli aspartate transcarbamoylase without regulatory subunits. Biochemistry 49: 7694–703

Mendes KR, Kantrowitz ER (2010b) The pathway of product release from the R state of aspartate transcarbamoylase. Journal of molecular biology 401: 940–8

Moreno-Morcillo M, Grande-Garcia A, Ruiz-Ramos A, Del Cano-Ochoa F, Boskovic J, Ramon-Maiques S (2017) Structural Insight into the Core of CAD, the Multifunctional Protein Leading De Novo Pyrimidine Biosynthesis. Structure (London, England: 1993) 25: 912–923 e5

Morris GM, Huey R, Lindstrom W, Sanner MF, Belew RK, Goodsell DS, Olson AJ (2009) AutoDock4 and AutoDockTools4: Automated docking with selective receptor flexibility. J Comput Chem 30: 2785–91

Newell JO, Schachman HK (1990) Amino acid substitutions which stabilize aspartate transcarbamoylase in the R state disrupt both homotropic and heterotropic effects. Biophys Chem 37: 183–96

Otwinowski Z, Minor W (1997) [20] Processing of X-ray diffraction data collected in oscillation mode. In Methods in Enzymology, pp 307–326. Academic Press

Pastra-Landis S, Foote J, Kantrowitz ER (1981) An improved colorimetric assay for aspartate and ornithine transcarbamylases. Analytical biochemistry 118: 358–363

Pastra-Landis SC, Evans DR, Lipscomb WN (1978) The effect of pH on the cooperative behavior of aspartate transcarbamylase from Escherichia coli. The Journal of biological chemistry 253: 4624–30

Ruiz-Ramos A, Lallous N, Grande-Garcia A, Ramon-Maiques S (2013) Expression, purification, crystallization and preliminary X-ray diffraction analysis of the aspartate transcarbamoylase domain of human CAD. Acta crystallographica Section F, Structural biology and crystallization communications 69: 1425–30

Ruiz-Ramos A, Velazquez-Campoy A, Grande-Garcia A, Moreno-Morcillo M, Ramon-Maiques S (2016) Structure and Functional Characterization of Human Aspartate Transcarbamoylase, the Target of the Anti-tumoral Drug PALA. Structure (London, England: 1993) 24: 1081–94

Serre V, Penverne B, Souciet JL, Potier S, Guy H, Evans D, Vicart P, Herve G (2004) Integrated allosteric regulation in the S. cerevisiae carbamylphosphate synthetase - aspartate transcarbamylase multifunctional protein. BMC Biochem 5: 6

Stebbins JW, Zhang Y, Kantrowitz ER (1990) Importance of residues Arg-167 and Gln-231 in both the allosteric and catalytic mechanisms of Escherichia coli aspartate transcarbamoylase. Biochemistry 29: 3821–7

Swyryd EA, Seaver SS, Stark GR (1974) N (phosphonacetyl) L aspartate, a potent transition state analog inhibitor of aspartate transcarbamylase, Blocks proliferation of mammalian cells in culture. Journal of Biological Chemistry 249: 6945–6950

Trott O, Olson AJ (2010) AutoDock Vina: improving the speed and accuracy of docking with a new scoring function, efficient optimization, and multithreading. J Comput Chem 31: 455–61

Tsuboi KK, Edmunds HN, Kwong LK (1977) Selective Inhibition of Pyrimidine Biosynthesis and Effect on Proliferative Growth of Colonic Cancer Cells. Cancer Research 37: 3080–3087

Wang J, Stieglitz KA, Cardia JP, Kantrowitz ER (2005) Structural basis for ordered substrate binding and cooperativity in aspartate transcarbamoylase. Proceedings of the National Academy of Sciences of the United States of America 102: 8881–6

Wang Q-S, Zhang K-H, Cui Y, Wang Z-J, Pan Q-Y, Liu K, Sun B, Zhou H, Li M-J, Xu Q, Xu C-Y, Yu F, He J-H (2018) Upgrade of macromolecular crystallography beamline BL17U1 at SSRF. Nuclear Science and Techniques 29: 68

West JM, Tsuruta H, Kantrowitz ER (2002) Stabilization of the R allosteric structure of Escherichia coli aspartate transcarbamoylase by disulfide bond formation. The Journal of biological chemistry 277: 47300–4

Wild JR, Loughrey-Chen SJ, Corder TS (1989) In the presence of CTP, UTP becomes an allosteric inhibitor of aspartate transcarbamoylase. Proceedings of the National Academy of Sciences of the United States of America 86: 46–50

